# Conformational Dynamics and Energy Landscapes of Ligand Binding in RyR1

**DOI:** 10.1101/167080

**Authors:** A. Dashti, D. Ben Hail, G. Mashayekhi, P. Schwander, A. des Georges, J. Frank, A. Ourmazd

**Affiliations:** Department of Physics, University of Wisconsin Milwaukee, 3135 N. Maryland Ave, Milwaukee, WI 53211, USA; Structural Biology Initiative, CUNY Advanced Science Research Center, City University of New York, New York, NY 10031, USA.; Department of Chemistry & Biochemistry, City College of New York, New York, NY 10031, USA.; Ph.D. Program in Biochemistry, The Graduate Center of the City University of New York, New York, NY 10016; Howard Hughes Medical Institute, Columbia University; Department of Biochemistry and Molecular Biophysics, Columbia University, 2-221 Black Building, 650 West 168^th^ Street, New York, NY 10032; Department of Biological Sciences, Columbia University

**Author notes:** These authors contributed equally (A. des Georges, J. Frank, A. Ourmazd). Corresponding authors: Corresponding authors’ contact details: Amedee des George; Joachim Frank; Abbas Ourmazd.

**Keywords:** Conformations of biological molecules, allosteric regulation, cryo-electron microscopy, manifold embedding, molecular movies

## Abstract

Using experimental single-particle cryo-EM snapshots of ryanodine receptor (RyR1), a Ca^2+^-channel involved in skeletal muscle excitation/contraction coupling, we present quantitative free-energy landscapes, reaction coordinates, and three-dimensional movies of the continuous conformational changes associated with the binding of activating ligands. Our results show multiple routes to ligand binding with comparable branching ratios. All high-probability routes involve significant conformational changes before and after the binding of ligands. We also present new insights into the local structural changes along the ligand-binding route, including accommodations at the calcium, ATP, and caffeine binding sites. These observations shed new light on the mechanisms and conformational routes to ligand binding.

---

The binding of a so-called ligand molecule to a macromolecular machine is the lynchpin of all functional and regulatory interactions in biology [1]. Despite sustained experimental and theoretical scrutiny for decades, the mechanisms by which ligands bind to macromolecular substrates [2, 3] remain the subject of intense debate (see, e.g., [1, 4, 5]). Broadly speaking, even the nature and sequence of events are controversial. In addition to fundamental scientific interest, these issues have important implications for drug design [6]. Experimentally determined multidimensional landscapes specifying the free energies of the conformations relevant to function offer a powerful framework for elucidating mechanisms [7, 8], but have remained largely inaccessible. Here, we present experimentally determined multidimensional energy landscapes of a macromolecular machine before and after the introduction of ligands. These landscapes reveal multiple routes to ligand binding with comparable branching ratios. Three-dimensional movies along these routes identify the continuous conformational changes involved in ligand binding at near atomic level, both overall and at specific binding sites. The approach can be used to elucidate the conformational changes associated with function in a wide range of systems. More generally, our work establishes that single-particle structural study of systems *in equilibrium* offers powerful insights into biological processes hitherto thought accessible only via *non-equilibrium*, time-resolved experiments.

We demonstrate the approach with reference to ryanodine receptor (RyR1), a fourfold-symmetric (C4) ion channel on the sarcoplasmic reticulum membrane. This macromolecule is a calcium-activated calcium channel critical to excitation/contraction coupling in skeletal muscle. Several recent cryo-EM studies have revealed in exquisite detail the many conformations assumed by the closed channel [9–11], with activation and gating of the channel upon ligand binding changing the nature of the conformational states [12]. But the plethora of discrete conformations revealed by powerful maximum likelihood classification methods [13, 14] provide scant guidance on the *sequence* of conformational changes involved in ligand binding and channel gating, the responsible *reaction coordinates*, the *energy landscapes* with and without ligands, or the *routes* to ligand binding and gating. Such information is essential for a deep understanding of these important processes, and the thermodynamics of function in general.

Using geometric machine learning techniques [15–17], we have previously demonstrated that experimental snapshots of single molecular machines idling in equilibrium can be used to determine their energy landscapes in terms of orthogonal coordinates describing *continuous* conformational changes relevant to translation [17]. Such landscapes reveal all conformations with energies up to an upper limit set by the vanishing occupation probability of high-energy states. The key point is that thermal fluctuations in equilibrium lead to sightings of all states up to the limit set by the number of snapshots in the dataset Supplementary section 1).

A key feature of the present study is the pooling of cryo-EM snapshots from two experiments. In one, RyR1 macromolecules were in equilibrium with a thermal bath. In the other, the macromolecules were in equilibrium with a thermal bath *and* a ligand reservoir [12] Supplementary section 2). This pooling of data has two important consequences. First, both species (with and without ligands, henceforth ±ligand) are described in terms of the same set of conformational reaction coordinates. Second, this approach reveals the heavily populated conformational conduits connecting the two species, thus identifying the lowest-energy routes relevant to ligand binding.

The approach outlined above allows us to determine the ±ligand energy landscapes for RyR1 in terms of the same binding-relevant reaction coordinates. Subject to reasonable assumptions, Fermi’s Golden Rule [18, 19] Supplementary section 3) is then used to estimate the transition probability between the two landscapes, with “hotspots” identifying the most probable transition points between the landscapes. Using the minimum-energy conduits on each landscape, we are thus able to compile, along high-probability routes, three-dimensional (3D) movies of ligand binding revealing changes in the overall channel conformation and at key binding sites. Our results demonstrate that the course of these continuous conformational changes differs in important ways from that obtained by morphing between discrete conformations [12].

The 791,956 cryo-EM snapshots of RyR1 molecules analyzed in this study comprised of about the same number of molecules in equilibrium with reservoirs with and without ligands (Ca^2+^, ATP, and caffeine) prior to cryo-freezing [12]. (For details see Supplementary section 2). These snapshots were grouped into 1,117 uniformly spaced orientational classes by standard procedures [20]. Geometric (manifold-based) analysis [17, 21] of the pooled dataset revealed at least four significant orthogonal conformational (reaction) coordinates, each describing a concerted set of continuous changes. Further detailed analysis was restricted to the first (i.e., the most important) two reaction coordinates (RC1 and RC2 for short). Conformational changes along RC1 involve the shell and pore, those along RC2 the activation core Supplementary section 4).

Fig. 1(A) shows the ±ligand energy landscapes of RyR1. Assuming the probability of a collision (not binding) with a ligand is independent of conformation, the probability of a transition between equivalent points on the two landscapes can be estimated from Fermi’s Golden Rule for the period immediately after the exposure of RyR1 molecules to the reservoir containing ligands Supplementary section 3). As shown in Fig. 1B, the inter-landscape transition probability displays specific “hotspot” regions, where a significant number of ligand-free and ligand-bound macromolecules display the same conformation. The most probable routes to ligand binding start from the region of lowest energy on the–ligand landscape (“START” in Fig. 1(A)), reach one of the hotspot transition points (“HOT”) with a probability of ~2%, cross to the +ligand landscape also with ~ 0.45% of the probability of a collision with a ligand, and terminate in the region of lowest energy on the +ligand landscape (“FINISH”). This means ~ 0.01% of collisions with a ligand lead to binding.

**Fig. 1.**
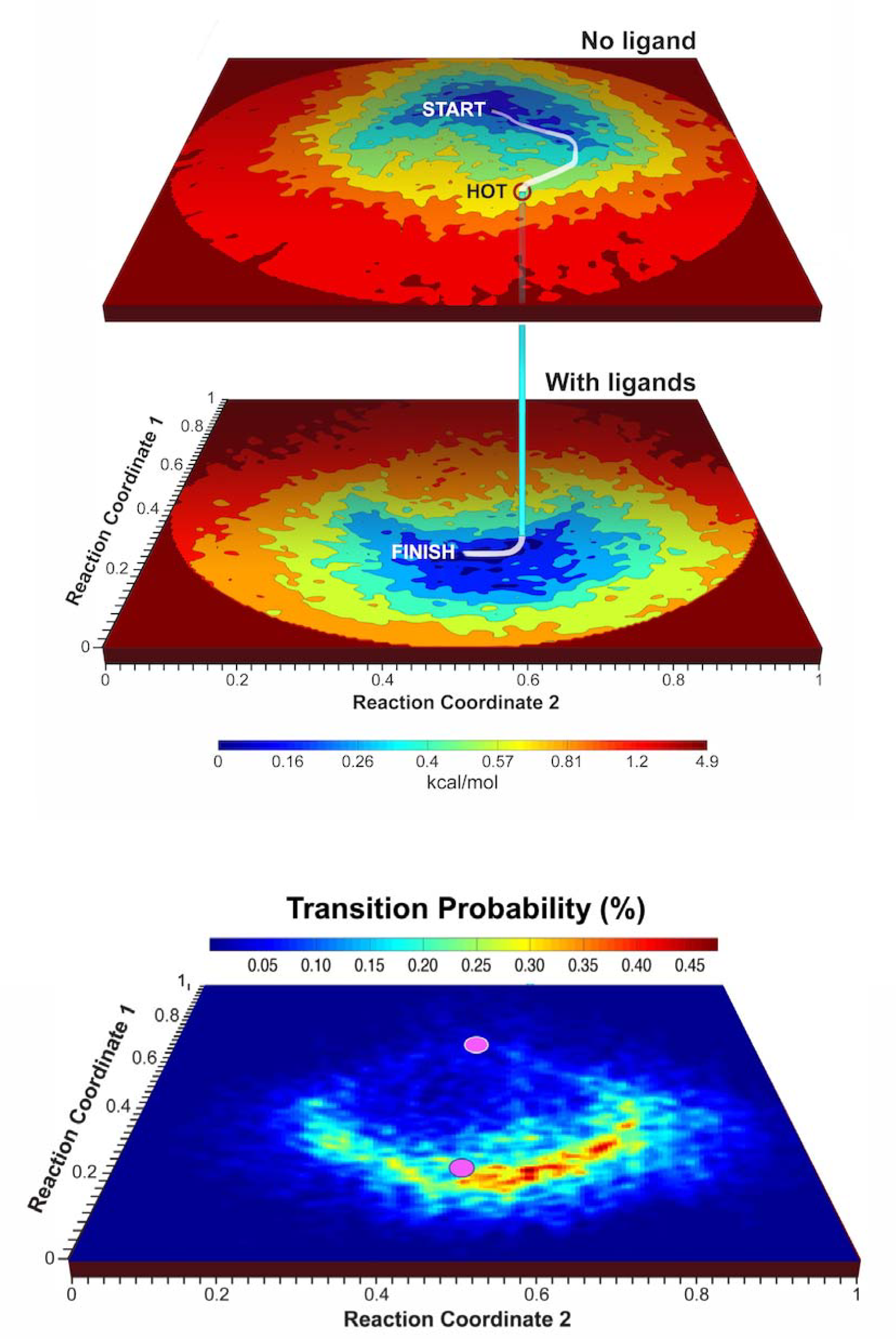
Energy landscapes for RyR1 with and without ligands, and map of transition probabilities to the with-ligand landscape. **(A)** Energy landscapes without, and with ligands (Ca^2+^, ATP, and caffeine), described in terms of the two most important orthogonal conformational reaction coordinates. The curved path represents a high-probability route to the binding of ligands. This path starts at the minimum-energy conformation of RyR1 without ligands (“START”), follows the conduit of lowest energy to a point with a high probability of transition to the with-ligands energy landscape (“HOT”), terminating at the minimum-energy conformation with ligands (“FINISH”). **(B)** Probability map for transitions from the energy landscape without ligands to the energy landscape with ligands. The axes are the same as in Fig. 1(A), with magenta discs indicating the positions of the minima of the energy landscapes at (0.7, 0.57) and (0.23, 0.56), respectively.

The displacement of inter-landscape transition hotspots (magenta in Fig. 1(B)) from minimum energy regions on both landscapes highlights the need for significant conformational changes before *and* after transition between the two energy landscapes. At the same time, the presence of several inter-landscape transition hotspots indicates a multiplicity of routes to ligand binding with comparable transition probabilities.

Fig. 2 shows the overall conformational evolution at selected points along the ligand-binding route shown in Fig. 1(A). (For estimates of the spatial resolution of our maps, and fitting and refinement of atomic coordinates, see Supplementary sections 5 and 6, respectively. For a 28-frame 3D movie of the conformational changes along this representative route, see Movies 1–4. For a discussion of these conformational changes, see Supplementary section 7. For a comparison of our results with those obtained via discrete classification by RELION [12, 14], see Supplementary section 8.)

**Fig. 2.**
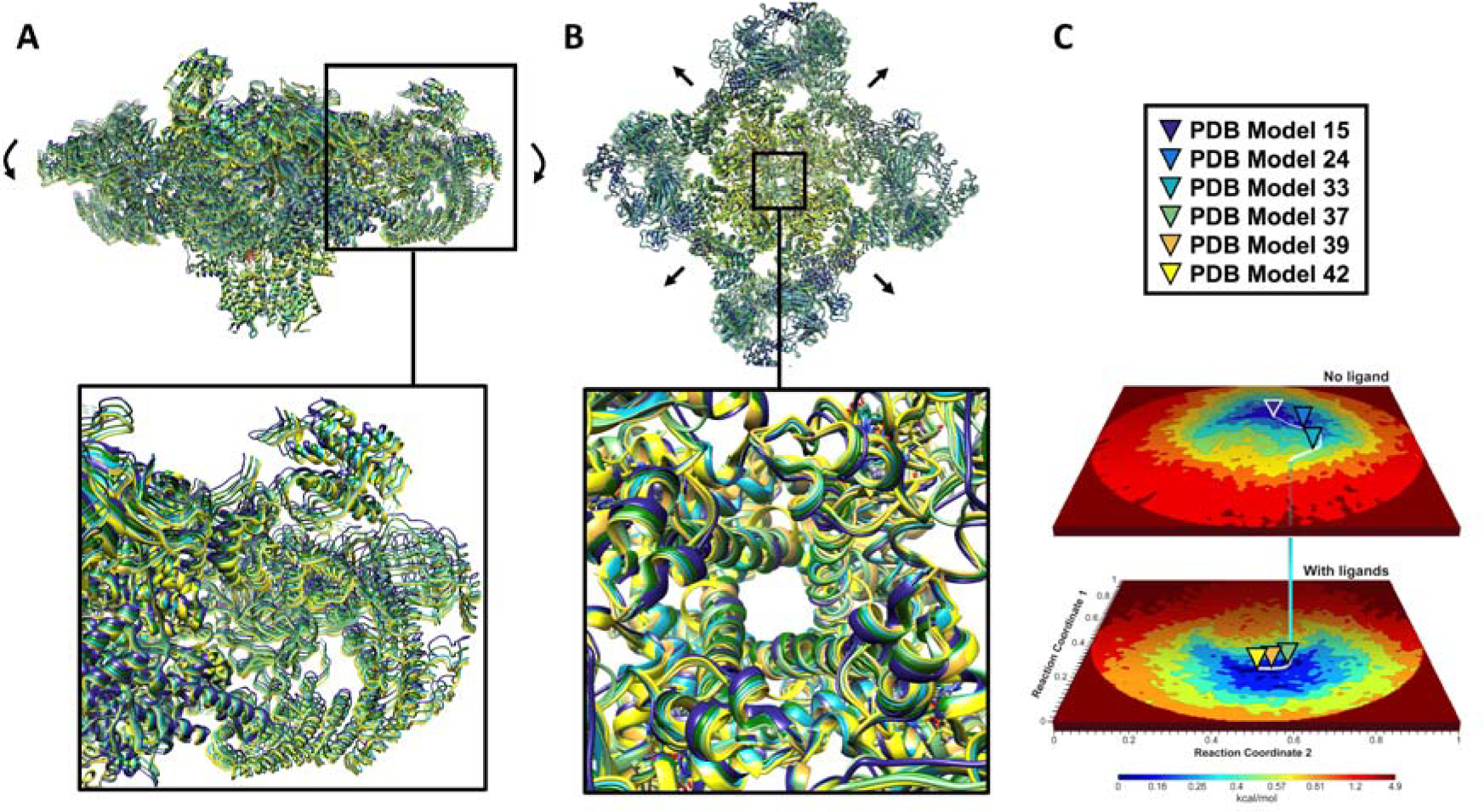
Five conformations of RyR1 along the ligand-binding route shown in Fig. 1. **(A)**, illustrating the overall conformational changes. The color of each conformation is the same as the triangle indicating its position on the trajectory. Each of the PDB model numbers corresponds to a structure deposited in the Protein Data Bank. Arrows indicate prominent movements. For 3D movies each consisting of 28 structures (frames), see Movies 1–4. **(A)** Trans-membrane view. **(B)** Cytoplasmic view.

Our results shed light on the longstanding question regarding the “population shift” vs. “induced fit” models. Broadly speaking, “population shift” [3] requires a conformational change *before*, “induced fit” [2] a conformational change *after* ligand binding. We find that ligand binding, at least in RyR1, proceeds via a continuum of conformations increasingly different from, and at higher energies than the minimum-energy conformation on the –ligand landscape. These higher-energy conformations are reached thermally via “population shift”. Collision with a ligand then transfers RyR1 to the +ligand energy landscape, where a downward slope in energy drives further continuous conformational changes to the minimum-energy, ligand-bound state. The conformational changes after collision with a ligand constitute an “induced shift”. Our results thus show that each of the two opposing models describes a different part of the actual process; at least for RyR1, binding entails specific conformational changes both before *and* after collision with a ligand.

The exact apportionment of the conformational changes before and after collision depends, of course, on the point at which the transition to the +ligand landscapes takes place. Although transitions can occur over a relatively broad region, the highest probabilities are concentrated at a few “hotspots” (Fig. 1(B)). The positions of the ±ligand energy minima relative to the broad region of significant transition probability indicate that most ligand-binding events in RyR1 involve a greater element of “population shift” than “induced fit”, as suggested in [22], but this could be system specific.

Our results also provide new insights into structural changes at specific binding sites (Fig. 3), and in the pore region (Fig. 4). A full discussion of these results and their implications for the gating mechanism of RyR1 is beyond the scope of the present paper. Here, we summarize a few salient points. First, the readily accessible conformations on the −ligand and +ligand energy landscapes overlap significantly. This indicates that, even in the absence of activating ligands (Ca^2+^, ATP and caffeine), RyR1 is able to assume, albeit with low probability, conformations previously thought to require the presence of a bound ligand [12, 23]. These conformational changes include, inter alia, changes in the activation and EF-hand domains, and partial pore opening Supplementary section 7, Movies 1–4). Second, moving along the ligand-binding conduit, we observe a motion of the cytoplasmic domain correlated with activation of the core and opening of the channel. The wing movements actuate a complex set of motions in the activation domain, which in turn bends the pore helix responsible for the opening of the pore Supplementary section 7). Third, the atomic coordinates of RyR1 obtained by modeling along the binding trajectory reveal the conformational changes at the binding sites of the ligands (Ca^2+^, ATP and caffeine) in the course of ligand binding, as outlined below.

**Fig. 3.**
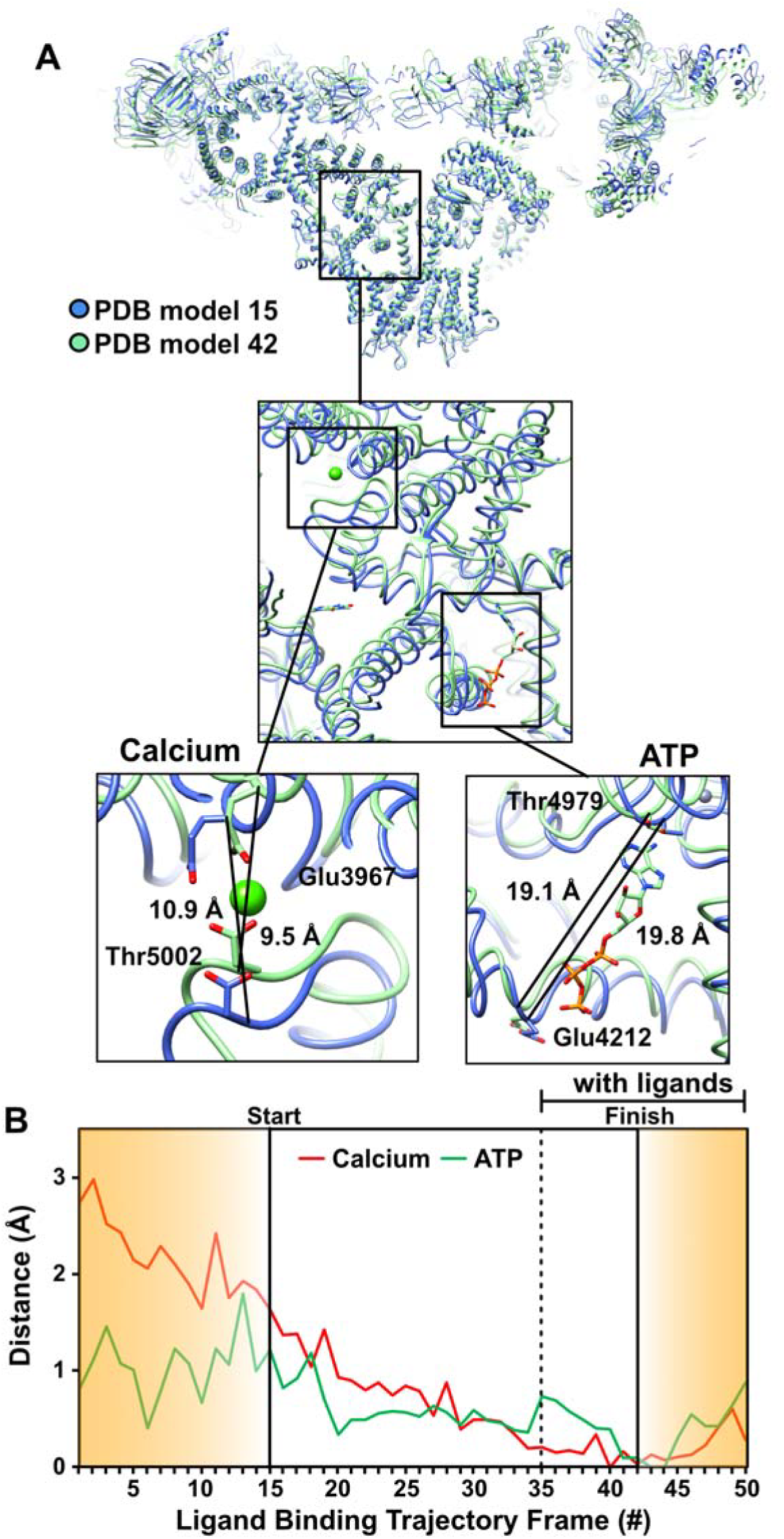
Conformational changes at RyR1 ligand binding sites along the route of Fig. 1. **(A)**, augmented with excursions from the minimum-energy points along RC1. These excursions are in the increasing RC1 direction on the no-ligand landscape, and in the decreasing RC1 direction on the with-ligand landscape. Each of the PDB model numbers corresponds to a structure deposited in the Protein Data Bank. **(A)** The general region, and the specific sites examined in detail. **(B)** Measured distances (in Å) between two amino acid backbones on opposite side of the ATP, and calcium, respectively. The amino acids used for measurement are represented in sticks, the rest of the molecule in blue ribbon for the model corresponding to “START” in Fig. 1(A), and in green ribbon for model corresponding to “FINISH”.

**Fig. 4.**
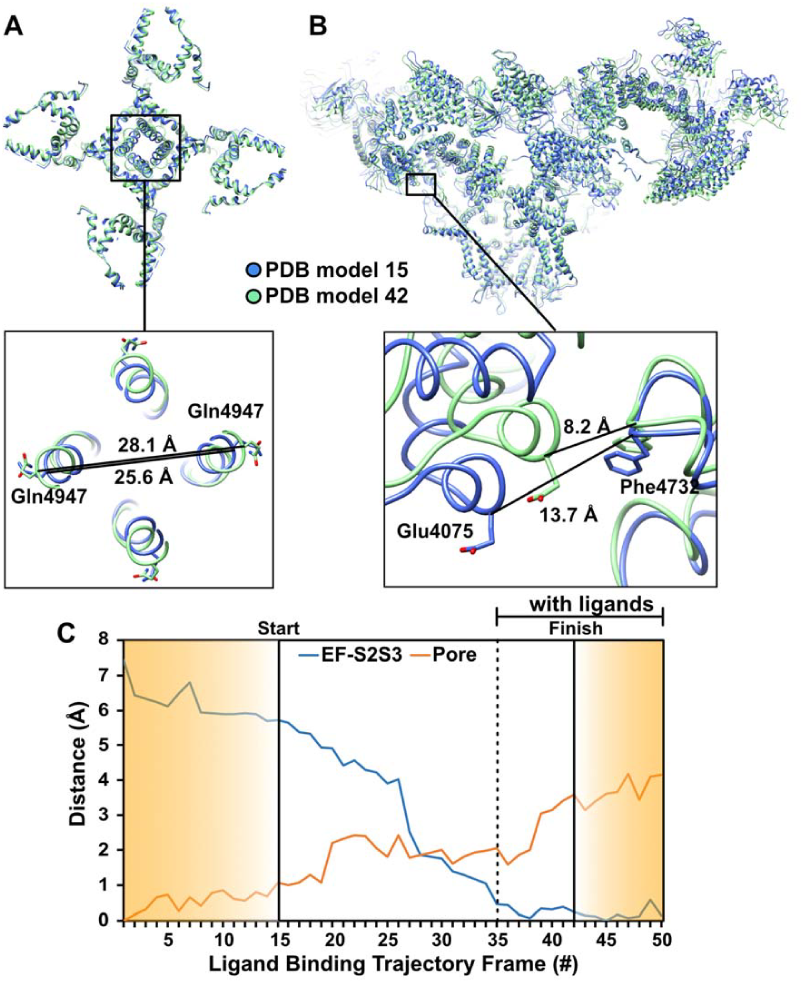
RyR1 conformational changes in the pore and EF-hand domains along the route of Fig. 1. (A), augmented with excursions from the minimum-energy points along RC1. These excursions are in the increasing RC1 direction on the no-ligand landscape, and in the decreasing RC1 direction on the with-ligand landscape. Distance variations (in Å) measured between Calpha backbone atoms of two opposing residues for the 50 frames along this trajectory. Amino acids used for measurement are represented as sticks, the rest of the molecule in blue ribbon for the model of frame #1 and in green ribbon for frame #50 at the two extremes of the trajectory. Each of the PDB model numbers corresponds to a structure deposited in the Protein Data Bank. **(A)** Pore opening measured at Gln4947; **(B)** Distance between EF-hand (EF) at Glu4075 and S2S3 domain at Phe4732. **(C)** Plot of Distance measurements for all 50 frames along the trajectory.

Starting at the minimum-energy point of the –ligand landscape, the Ca^2+^ binding site gradually contracts until the transition to the ligand-bound landscape is effected, after which the binding-site conformation stabilizes in its ligand-bound state (Fig. 3). In contrast, the ATP binding site does not display a large distance change along the entire conduit connecting the energy minima of the ±ligand landscapes (Fig. 3). This binding site thus seems to require little, if any accommodation to bind its ligand, indicating the ATP-bound conformation of the domain is close to the minimum energy conformation, whether ATP is bound or not. This is expected from the constitutively bound nature of ATP at physiological concentrations [12, 24]; ATP is not a physiologically activating ligand. ATP’s main role appears to involve rigidifying the pore-helix/C-terminal domain hinge, thus performing a structural role necessary for gating.

We now discuss the more general implications of our approach. First, biological function involves a rich set of *continuous* conformational changes inadequately described by discrete classes of unknown relationship. Specifically, movies obtained by morphing between discrete clusters may involve conformational routes not central, or even relevant to the function of interest. Second, *thermal fluctuations in equilibrium* lead to sightings of all states up to a limit set by the number of snapshots in the dataset. This allows one to compile the energy landscapes needed for a rigorous description of the thermodynamics of function. And third, the course of a biological *process* can be inferred by pooling data from ensembles in equilibrium with reservoirs corresponding to the initial and final states of the process. Demonstrating the ability of this approach to provide new insights into biological function has been an overarching goal of this paper. We believe the energy landscapes, the inter-landscape transition maps, and the new information on the nature and sequence of events needed for ligand binding offer important new insights into longstanding questions on ligand binding, and offer a general framework for understanding macromolecular function in general.

Clearly, equilibrium measurements cannot answer all questions regarding dynamics. It would thus be illuminating to compare the present results with those obtained from observation of non-equilibrium ensembles engaged in reaction. Also, using larger equilibrium datasets, it would be interesting to investigate the role of conformations lying at higher energies. These constitute future tasks.

## Conclusions

We have presented a new approach to determining the structural and thermodynamic factors underlying a wide range of biological processes, and demonstrated its power with reference to ligand binding in RyR1, the calcium release channel in skeletal muscle. Our results reveal multiple routes to ligand binding in RyR1, and elucidate the mechanisms underlying in this vital process. The approach is general, and thus applicable to a wide range of systems and processes.

## Acknowledgments

We acknowledge valuable discussions with E. Lattman, G. Phillips, M. Schmidt, and members of the UWM data analysis group. The research conducted at UWM was supported by the US Department of Energy, Office of Science, Basic Energy Sciences under award DE-SC0002164 (algorithm design and development, and data analysis), and by the US National Science Foundation under awards STC 1231306 (numerical trial models) and 1551489 (underlying analytical models). The work performed by JF was supported by HHMI, NIH GM55440, and NIH GM29169. The work performed by DBH and AG was supported by CUNY.

## Supplementary Materials

**Movies 1, 2, 3, 4**

## Author contributions

AD designed and implemented the data-analytical approach, a robust pipeline for driving energy landscape, transition probability maps and continuous conformational changes from Cryo-EM snapshots.

AG processed the cryo-EM data using standard methods. AG and DBH designed the molecular modeling and distance measurement strategy. DBH performed the molecular modeling and distance measurements.

GM helped with implementation of data-analysis algorithms, mainly for the energy landscapes and the continuous conformational movies.

PS co-developed the geometric manifold-based approach, co-investigated the effect of coarse graining on energy landscapes, and helped implement the software.

JF co-designed the study, analyzed the results in terms of the concepts of single-particle cryo-EM and the dynamics of molecular machines, and co-wrote the paper.

AO co-designed the study and the manifold-based data-analytical algorithms, identified and co-analyzed the effect of coarse-graining, proposed the ligand-binding model and formalism for estimating interlandscape transition probabilities, and wrote the paper.

AO, AD, AG, DBH and JF analyzed and interpreted the results.

